# Examining Selection Dynamics and Limitations in Multi-round Protein Selection of High Diversity Libraries

**DOI:** 10.1101/2025.11.09.687419

**Authors:** John Z. Chen, Barnabas Gall, Tommy Y. Lu, Isabella Heslop, Daniel Hesselson, Christoph Nitsche, Wai-Hong Tham, Richard J. Payne, Colin J. Jackson

## Abstract

Proteins and peptides underpin essential biological functions and technological applications, from targeting disease-relevant interactions to providing broad enzymatic activities. However, engineering molecules with desired properties remains difficult, owing to complex sequence-structure-function relationships and the lack of data on specific systems. Experimental selection strategies, including directed evolution, phage display, and mRNA display, address this challenge by leveraging high diversity libraries and iterative enrichment under defined selection pressures. This allows for the identification of candidates without requiring extensive prior knowledge, and can generate extensive datasets for use in machine learning. While many selection systems exist, comparisons across different selection approaches are hindered by the lack of a unifying analytical framework. Here, we present a set of broadly applicable analyses for assessing selection dynamics in multi-round or multi-condition experiments, ranging from position level analysis of sequence properties to full sequence space mappings through protein language model embeddings. Using the toolset to analyze a variety of different datasets in parallel, we explore the potential effects of diversity, coverage, and reproducibility, offering generalizable insights to guide experimental design, interpretation, and troubleshooting across protein and peptide discovery platforms.

## Introduction

Proteins and peptides are central to virtually all biological systems, performing a vast range of functions that make them attractive candidates for diverse applications. For example, their critical role in protein–protein interactions (PPIs) underlie essential cellular processes such as signal transduction, immune responses, and pathogen invasion (Lu *et al*., 2020; Greenblatt *et al*., 2024). Dysregulation of PPIs is implicated in many diseases, and targeting specific interactions with engineered peptides or antibodies/nanobodies has emerged as a promising strategy for therapeutic and diagnostic development, as demonstrated in cases like malaria (Wright *et al*., 2014; Dietrich *et al*., 2022) and SARS-CoV-2 (Norman *et al*., 2021; Johansen-Leete *et al*., 2022; Thijssen *et al*., 2023). Similarly, design of enzymes with tailored activities, such as those capable of degrading synthetic plastics (Tournier *et al*., 2020; Joho *et al*., 2023), highlight the potential of proteins in solving pressing technological and environmental challenges. Despite this promise, discovering or designing proteins and peptides with precisely tuned properties remains challenging due to the complexity of their structure–function relationships. Recent advances in machine learning have enabled models to learn patterns from large protein datasets, allowing for predictions of general features like structure or function (Madani *et al*., 2020; Rives *et al*., 2021; Hayes *et al*., 2025; Yuan *et al*., 2025). However, these models are challenging to construct in specific systems where training data is sparse or non-existent (Dobbelaere *et al*., 2021; Yuan *et al*., 2025), limiting their effectiveness in guiding protein engineering in niche or novel contexts.

In the absence of prior knowledge, selection from diverse peptide or protein pools is a powerful strategy for discovering candidates with desired properties, such as novel or enhanced activity. Directed evolution exemplifies this approach by subjecting a starting protein to iterative rounds of mutation and selection, incrementally navigating the mutational landscape toward improved function (Yuan *et al*., 2005; Bloom and Arnold, 2009; Tokuriki *et al*., 2012; Alvizo *et al*., 2014; Yang *et al*., 2019). In contrast, display technologies like phage display (Omidfar and Daneshpour, 2015; Saw and Song, 2019; Zhang, 2023) and mRNA display, e.g., the Random non-standard Peptide Integrated Discovery (RaPID) system (Passioura and Suga, 2017; Goto and Suga, 2021), begin with vast, highly diverse libraries and progressively enrich for functional sequences without needing to introduce new mutations during selection. Regardless of the method, two factors are essential for success: the availability of sufficient mutational diversity and the application of a specific selection pressure (Yuan *et al*., 2005; Otto and Whitlock, 2013). The advent of high-throughput sequencing has allowed these approaches to build comprehensive profiles of library diversity and track the enrichment of specific variants across selection rounds (Thijssen *et al*., 2023; Ullrich *et al*., 2024; Cole *et al*., 2025). This allows researchers to assess the full spectrum of sequences and their relative performance, reducing reliance on random sampling and improving hit identification efficiency. Moreover, the resulting sequence–function data provides insights that can inform rational design strategies, or be integrated into hybrid approaches where mutagenesis in subsequent rounds is guided by observed trends in previous selections (Yang *et al*., 2019; Minot and Reddy, 2024; Thomas *et al*., 2025).

While advances in peptide and protein selection and design have generated a wealth of studies and diverse outcomes, there remains a lack of a common framework for comparing results across different selection methods. In this work, we present a set of common analyses to examine selection dynamics in multi-round or multi-condition experiments using high-diversity libraries. The toolset provides a broad overview of a dataset, including sequencing coverage, round to round correlations, plots to highlight positional sequence properties and a full sequence space visualization using protein language model embeddings. These methods are broadly applicable to data from various platforms, including mRNA display, phage display, iterative design, and directed evolution approaches. We highlight key considerations such as library diversity, variant coverage, and reproducibility, all of which can significantly influence the outcome of selection experiments. These generalizable analyses can help experimenters assess the progression and quality of their selections, and identify cases where further selection or troubleshooting may be warranted.

## Results

### Generalizable approach for analyzing multi-round selection data

To examine the selection dynamics in high diversity libraries, we employ a suite of analyses that examine different aspects of the selection outcomes (**Fig. 1**). We assess the distribution of variant frequencies across rounds to determine whether selection is acting to enrich variants of interest, since there should be increasing frequencies of a subset of sequences along with depletion of other sequences. We also look at correlation across consecutive rounds of selection, to see if the selection is behaving consistently throughout the course of the experiment; if consecutive rounds are correlated, it indicates consistent selection toward a common subpopulation. After the final rounds of the experiment, we look at the per position amino acid distribution, as well as biophysical properties for any common patterns. The presence of preferred residues or motifs indicates the selected subpopulations share common patterns; variation in a subset of the positions could also highlight mutational flexibility. Finally, we visualize the shift of the population via dimensionality reduction (UMAP) of protein language model (pLM) embeddings, allowing each sequence to be represented as a point in 2D space. This allows for visual inspection of the selection process at a global level and can highlight the nature of the selected subpopulation such as the number of “hotspots” of enrichment, and differences in selection endpoints between conditions. As we will show, these methods are generalizable to library selection data of different origins with small modifications.

**Figure 1.**
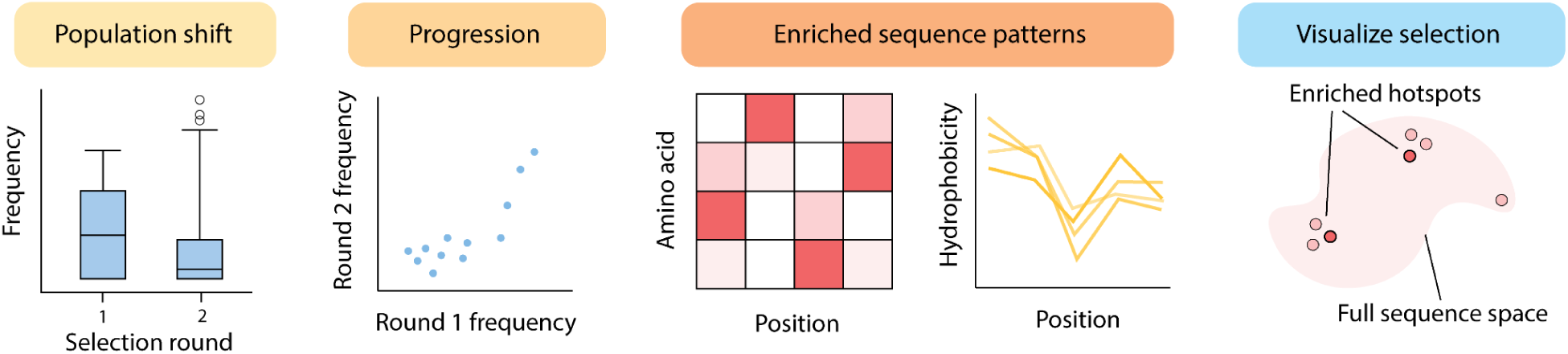
Overview of analyses to examine multiround selection data. Once a dataset of sequences and mutations has been obtained from deep sequencing, these analyses can be used to gain a quick and intuitive overview of the data. Overall effectiveness of selection can be monitored by shifts in population frequencies. The progression of selection can be examined by comparing frequencies between rounds. The sequence patterns at a given round can be summarized by their amino acid composition, or biophysical parameters. The full sequence space can be visualized with a UMAP plot of sequences based on their protein language model embeddings.

### Case 1: mRNA display selection of cyclic peptides against SARS-CoV-2 spike protein

In this study, Thijssen *et. al.* performed mRNA display on two libraries of 15 NNK randomized codons (differing by D- or L-Tyr as the initiator) to select for binding to SARS-CoV-2 spike ectodomains (Thijssen *et al*., 2023). We analyzed the part of the experiment directly targeting the spike protein with selection and sequencing over 6 rounds. After filtering for sequences with minimum Phred scores ≥ 20, read counts per sample were fairly low at ∼10k reads per sample, but nonetheless led to clear patterns of selection. The per-sequence frequency after each round of selection shows a shift of the distributions’ tails toward higher frequency, while the bulk of sequences accumulate at lower frequencies (Fig. 2a). This shows selection is effective at purging most sequences, while enriching a smaller subset. As the rounds of selection progress, we also see greater sequence overlap and better correlation between successive rounds (Fig. 2b), showing the selection is stabilizing toward a certain sequence subpopulation. When viewing the sequence patterns at round 6, specific amino acid preferences for each position in the randomized region are apparent (Fig. 2c), with a stronger pattern for the L-Tyr initiated library. This is also seen in the biophysical properties (Fig. 2d), where top hits for L-Tyr have more consistent hydrophobicity and accessible surface area profiles compared to D-Tyr. Indeed, the most effective inhibitors were from the L-Tyr library (Thijssen *et al*., 2023). Finally, viewing the entire selection sequence space as a UMAP of the pLM embeddings, it is clear that the D- and L-Tyr libraries are selected toward different parts of sequence space, eventually converging on several high frequency hotspots. Overall, the SARS-CoV-2 spike selection demonstrates consistent selection toward a common sequence pattern for each library.

**Figure 2.**
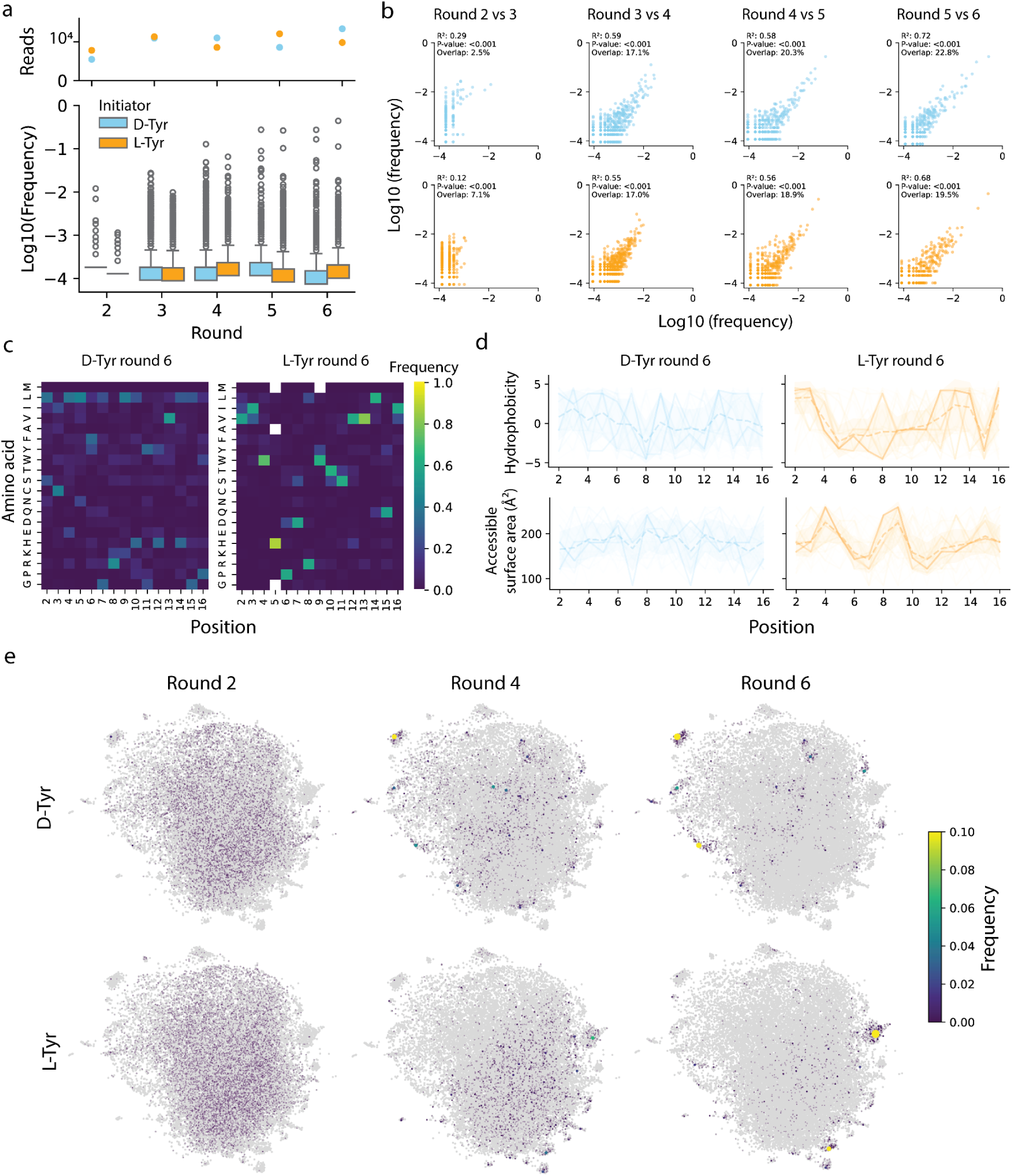
mRNA display of cyclic peptides against SARS-CoV-2 spike protein. (**a**) Total number of reads and sequence frequency after each round of selection. Peptides with D- and L-Tyr initiators are separated by color, maintained throughout the figure. (**b**) Correlation of sequence frequencies between consecutive rounds of selection. The R2 and P-value are shown for a linear fit. The overlap between rounds is calculated as sequences shared/total unique. (**c**) Heatmap of positional frequencies for each amino acid. (**d**) Plot of top 100 most frequent sequences as line plots by biophysical property. The dashed line is the positional average and the envelope is the position standard deviation. (**e**) UMAP display of sequences over the course of selection. All encountered sequences are in grey, while sequences present in the current round are colored and have point sizes scaled by frequency.

A strength of high-throughput screens is the ability to explore a vast diversity of potential binding solutions for a given target. A randomized library (NNK/S) of 15 codons, as commonly used in RaPID peptide libraries amounts to a maximum theoretical diversity of 6.8 x 10^19^ amino acid variants, or 3.8 x 10^22^ codon variants, as calculated using Peptider (Sieber *et al*., 2015) (**Table 1**). RaPID experiments have throughput for ∼10^12^ sequences in the initial mRNA display (Goto and Suga, 2021), where such a library would have only ∼10^-8^ (1 in 100 million) coverage of its full diversity. Any repeated sampling would likely produce completely different results given the sheer size of the sequence space. In this study, two candidates of the top 15 tested were found capable of pseudoviral neutralization, giving a conservative estimate of at least 1 hit in 5×10^11^ such randomized sequences. Hence, covering a large search space is required and smaller sample sizes are more likely to miss such hits. Although the hits seem rare, there are potentially 10^8^ times more hits encoded in the full-sized randomized library (assuming completely random distribution) and there are potentially billions of binding solutions in this sequence space for any given binding function.

**Table 1.**
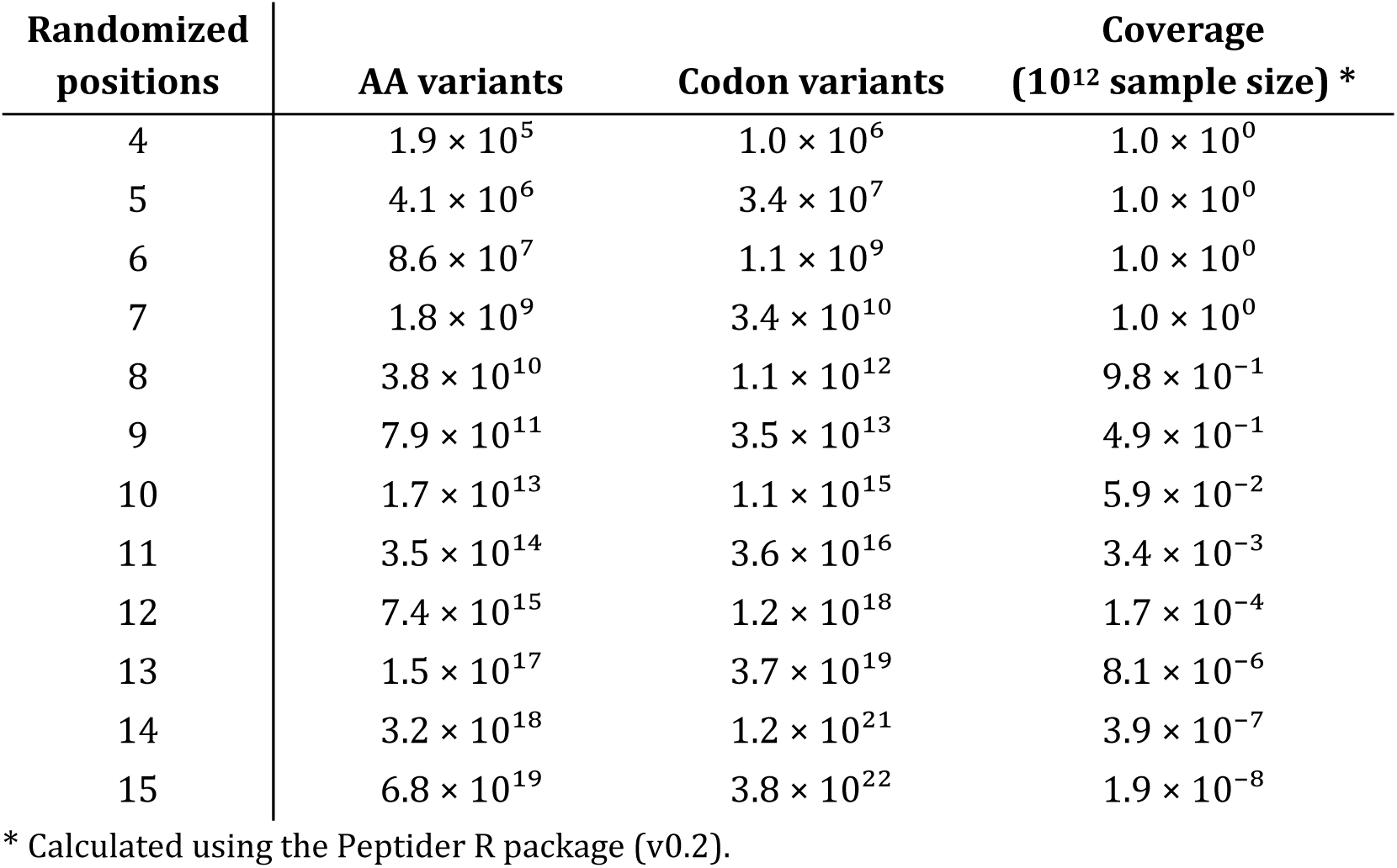
Estimated coverage of randomized (NNK/S) peptide libraries of a given length.

### Case 2: mRNA display selection cyclic peptides against main protease of SARS-CoV-2

In this study by Johansen-Leete *et. al.*, RaPID mRNA display was performed against the main protease (M^Pro^) of SARS-CoV-2 (Johansen-Leete *et al*., 2022). Two randomized libraries (differing by D- or L-Tyr as the initiator) of 4-15 NNS codons were mixed proportionately into a single library. The library then underwent 9 rounds of selection with sequencing performed at rounds 7 and 9. The original study noted a lack of selection at round 7, prompting further selection to round 9 which also did not show conserved patterns. Here we examine the selection in detail. The per-sequence frequencies of the libraries remain almost unchanged between rounds 7 and 9 (Fig. 3a), both remaining below 1% (10^-2^) even at the highest frequencies with no upward trend. The degree of frequency change for M^pro^ at round 9 is similar to the beginning of selection for the spike selection (Fig. 2a, round 2). The correlation between rounds 7 and 9 is quite weak, suggesting no consistent enrichment has occurred (Fig. 3b**, Supp.** Fig. 1 for separate lengths). When looking at the per position amino acid frequencies, again no clear pattern emerges (Fig. 3c), with weak bands of non-polar amino acids (Leu, Val) across most positions, which suggests the peptide pool is overall retaining hydrophobic “sticky” peptides. The lack of pattern is shared for the biophysical properties of the top 100 sequences (Fig. 3d). Finally, examining the sequence space, there are no distinct changes between rounds and sequences are scattered across the space with no notable hotspots (Fig. 3e).

**Figure 3.**
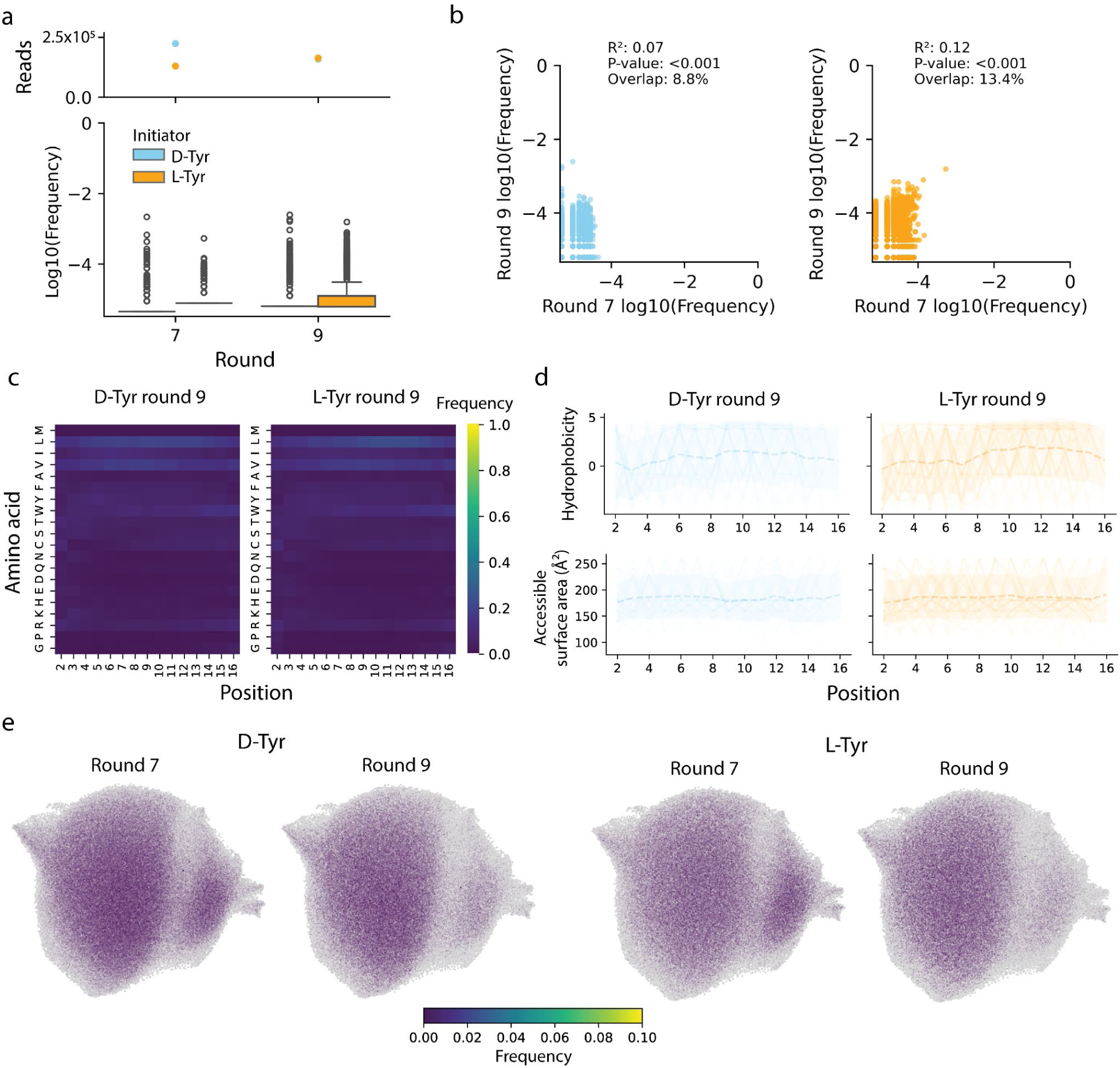
mRNA display of cyclic peptides against the main protease of SARS-CoV-2. (**a**) Total number of reads and sequence frequency after each round of selection. Peptides with D- and L-Tyr initiators are separated by color, maintained throughout the figure. (**b**) Correlation of sequence frequencies between rounds 7 and 9 of selection. The R2 and P-value are shown for a linear fit. The overlap between rounds is calculated as sequences shared/total unique. (**c**) Heatmap of positional frequencies for each amino acid. All lengths in the library are plotted together with the linker region removed. Heatmaps of separate lengths can be found in **Supp.** Fig. 1. (**d**) Plot of top 100 most frequent sequences as line plots by biophysical property. The dashed line is the positional average and the envelope is the position standard deviation. (**e**) UMAP display of sequences over the course of selection. All encountered sequences are in grey, while sequences present in the current round are colored and have point sizes scaled by frequency.

In the original study, the lack of consensus after selection was attributed to the structural/functional heterogeneity in the system. As M^Pro^ is catalytically active as a dimer, cross-linking was required to recapitulate the catalytic form. However, the cross-linking was incomplete (Fig. 2b in Johansen-Leet *et. al.* 2022) and the presence of the monomer could have added heterogeneity (Johansen-Leete *et al*., 2022). To investigate other possible causes, we considered the possibility that the study’s library design of pooling multiple peptide lengths may have diluted potential hits. However, the number of codon variants added by the varying lengths (4-14 codons, added proportionally) is only ∼3% that of the 15 codon library (**Table 1**) and this is unlikely to be significant. The sequence patterns in the final round show Leu, Arg and Val as the most common across positions (Fig. 3c). These are also among the most common in the NNK codon set as Leu and Arg have 6 codons, and Val has 4 codons. This suggests that selection strength was also weak regardless of target heterogeneity, as the library as a whole appears to have been only slightly enriched from randomized frequencies. Despite this, the study was able to identify high affinity binders with inhibitory activity (best hit with 70nM IC_50_) against the SARS-CoV-2 M^Pro^ (Johansen-Leete *et al*., 2022), which highlights the power of large library screening approaches.

### Case 3: Phage display selection of nanobodies targeting *Plasmodium falciparum* Pfs230

In recent work by Dietrich *et. al.*, an alpaca was immunized with the first two 6-cys domains of Pfs230 (Pfs230 D1D2), a malaria surface protein essential for fertilization and gamete fusion during malaria sexual reproduction, (Dietrich *et al*., 2022) to generate a 10^6^ nanobody phage display library, which was then selected over two rounds for binding to Pfs230, both of which were sequenced to a depth of ∼1M reads after filtering. Although the library was only selected over two rounds, there were clear signs of strong selection, as the frequencies of the variants in the library showed a substantial shift between rounds 1 and 2 (Fig. 4a). Some variants were already observed at very high frequencies after round 1 (top hit occupies ∼10% of the population), and there was further depletion of low frequency variants in round 2. Comparing selection between rounds, there is consistent behavior for high frequency variants (present at > 1% of the population), while lower frequency variants are less correlated (Fig. 4b). To identify sequence patterns, we focus on the complementary-determining region 3 (CDR3), which is the most varied and most likely responsible for differences in binding (Dietrich *et al*., 2022). When aligned to the longest CDR3 (25aa) with the highest frequency, we see patterns in the CDR3 that become stronger across rounds (Fig. 4c), which is also reflected in the biophysical properties (Fig. 4d). There appears to be distinct groups of long and short CDR3s (as opposed to a gradual range of lengths) (Fig. 4c), which is reflected in the final hits in the study (Dietrich *et al*., 2022). Examining the sequence space, hotspots are already present after the first round (Fig. 4e), and in round 2 there is further shift away from low frequency regions across the background toward these hotspots.

**Figure 4.**
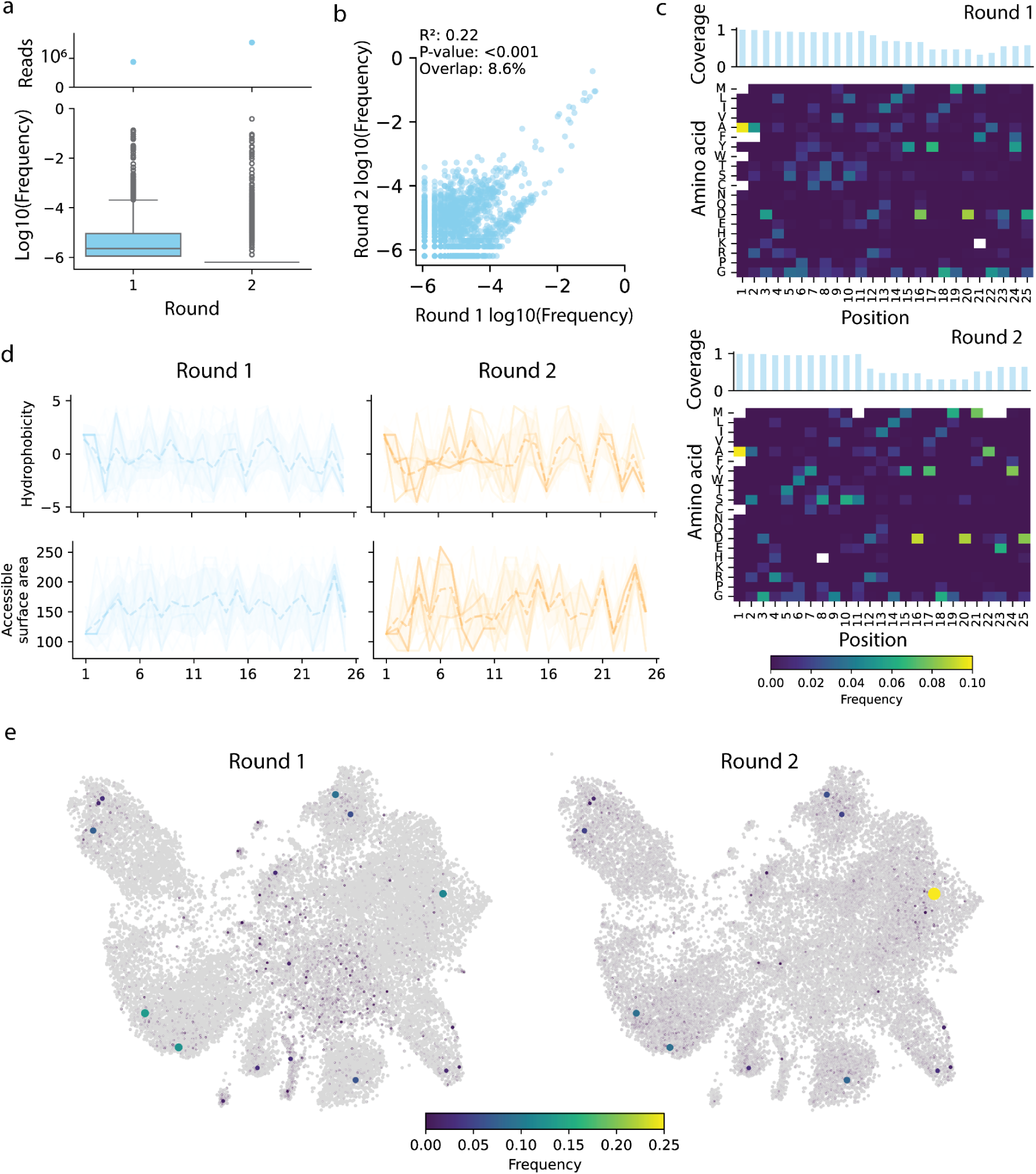
Selection in CDR3 region of nanobodies targeting *Plasmodium falciparum* Pfs230. (**a**) Total number of reads and sequence frequency after each round of selection. (**b**) Correlation of sequence frequencies between rounds 1 and 2 of selection. The R^2^ and P-value are shown for a linear fit. The overlap between rounds is calculated as sequences shared/total unique. (**c**) Heatmap of positional frequencies for each amino acid, aligned to the longest CDR3. Coverage of the aligned position is shown at the top. (**d**) Plot of top 100 most frequent sequences as line plots by biophysical property. Dashed line is the positional average and the envelope is the position standard deviation. (**e**) UMAP display of sequences over the course of selection. All encountered sequences are in grey, while sequences present in the current round are colored and have point sizes scaled by frequency.

This study of nanobody phage display showcased strong selection and isolation of the desired function in just two rounds of selection, compared to a higher number typically required by mRNA display. Of the 12 nanobodies selected for characterization, 11 had nanomolar affinities (0.45-16.15 nM) (Dietrich *et al*., 2022). Notably, this is in spite of the library size being orders of magnitude smaller (10^6^ in this study vs 10^12^ for mRNA display). One possibility is that many more solutions are present in the mRNA display experiments but remain unknown due to lack of characterization, but this is perhaps unlikely on the order of 10^6^ when the top hits are not always consistent. Rather, the enhanced efficacy of this screen is likely due to prior enrichment and selection. The alpaca was immunized specifically with the antigen before library generation, enriching nanobodies with affinity to the antigen. Furthermore, nanobodies in general have already been selected in nature to be protein binders. This would decrease the amount of further selection needed to find a satisfactory hit compared to starting from a fully randomized, unselected sequence space. There is therefore a trade-off to be considered in selecting binders using a pre-selected system adapted from nature, or an unselected synthetic library that could yield completely novel binding solutions.

### Case 4: Phage display selection of bismuth bicyclic peptides against streptavidin

Recent work by Ullrich et. al. utilized a semi-randomized library with the peptide motif ACX_4_CX_4_C (Ullrich *et al*., 2024). These peptides are linear, but are made bicyclic by coordinating to metal (Bi^3+^) or metalloid (As^3+^) ions. In this experiment, four rounds of selection against streptavidin were performed and only the final round was sequenced. Unlike the other examples, this study features the same peptide library, selected in the presence of different ions (BiBr_3_ and gastrodenol, a.k.a. bismuth tripotassium dicitrate, for Bi^3+^, and NaAsO_2_ for As^3+^); the NaAsO_2_ condition was conducted in duplicate. Hence, we can use our computational pipeline to compare different selection conditions, as opposed to multiple rounds under the same condition.

The distribution of peptide frequencies after 4 rounds of selection are comparable between conditions, with each having achieved high frequency hits (>1%), and the majority of peptides having depleted to low frequencies (Fig. 5a). We can see both Bi containing conditions correlate better with each other (R^2^=0.52) than with the As condition (R^2^ ≤ 0.14) (Fig. 5b), indicating consistent selection. The As condition replicates self-correlate to a similar degree (R^2^=0.54) as the two Bi conditions. Within the Bi conditions, BiBr_3_ shows multiple enriched residues at each position (Fig. 5c), while gastrodenol exhibits a weaker but similar pattern. The first X_4_ region of the two NaAsO_2_ replicates show a preference for HPQN, but are more varied in the second region (Fig. 5c). The pattern is less distinct in terms of biophysical properties by position (Fig. 5d). As noted in the original study, patterns expected to interact with streptavidin (amino acid motif HPQ/M) appear frequently, (Ullrich *et al*., 2024). In sequence space, we find distinct hotspots shared between each metal group, showing different conditions lead to different selection outcomes in sequence space (Fig. 5e). The sequence space visualization also makes other observations more intuitive;e.g., the weaker enrichment in the gastrodenol selection is due to more peptide backgrounds with intermediate frequencies (near depleted, but higher than background of other conditions) (Fig. 5a). The replicates of the As condition have overlapping hotspots but different frequencies, which explains the limited replicate correlation R^2^ of 0.54.

**Figure 5.**
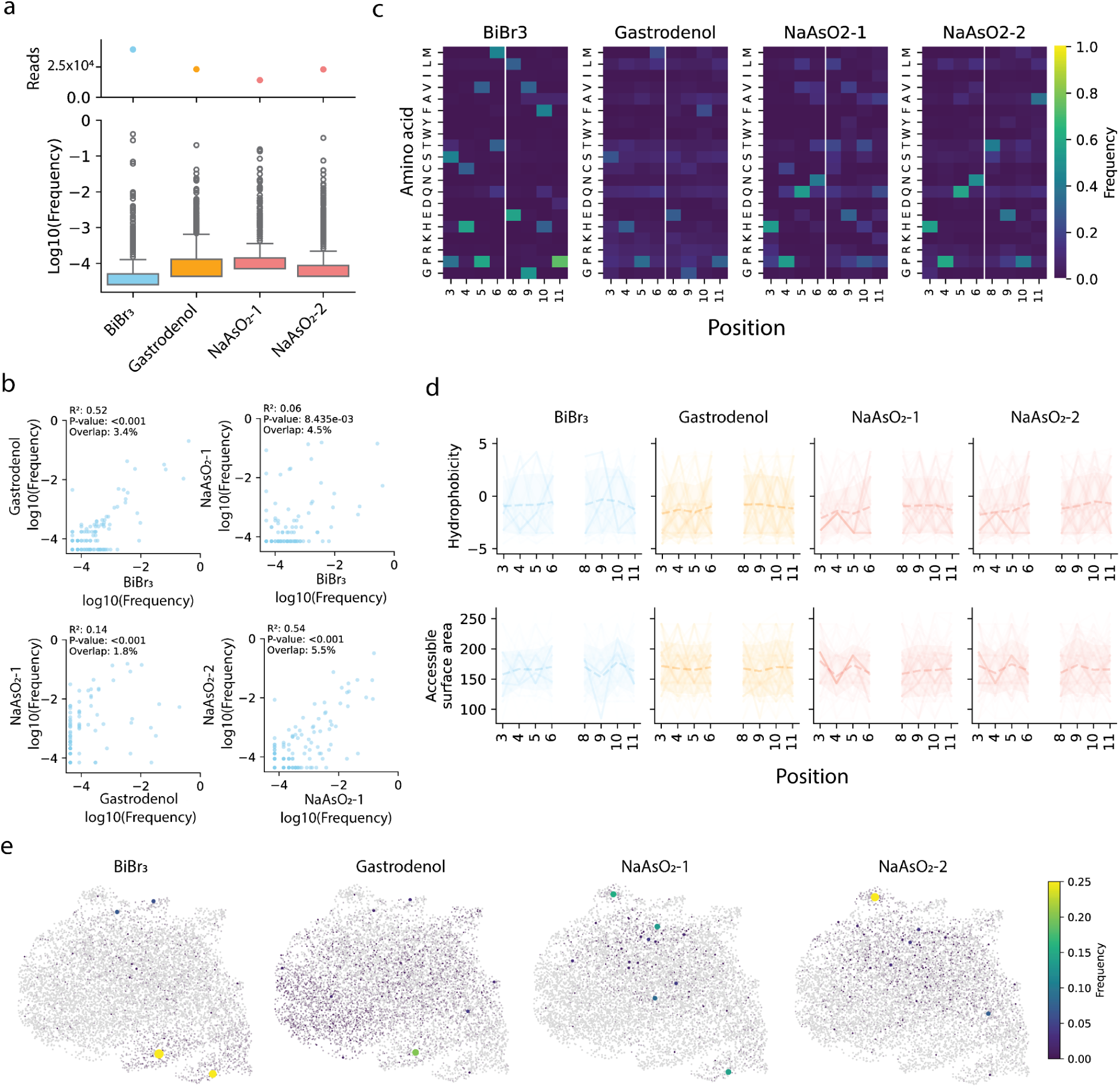
Bicyclic peptides selected against streptavidin under different metal conditions. (**a**) Total number of reads and sequence frequency after round 4 of selection for all conditions. The NaAsO2 condition was performed in duplicate. (**b**) Correlation of sequence frequencies between each type of metal selection. The R2 and P-value are shown for a linear fit. The overlap between rounds is calculated as sequences shared/total unique. (**c**) Heatmap of positional frequencies for each amino acid. (**d**) Plot of top 100 most frequent sequences as line plots by biophysical property. The dashed line is the positional average and the envelope is the position standard deviation. (**e**) UMAP display of sequences over the course of selection. All encountered sequences are in grey, while sequences present in the current round are colored and have point sizes scaled by frequency.

This study includes the same library of sequences selected under different conditions and in duplicate. While the two Bi conditions showed agreement, it is interesting that they do not align completely (some overlap in top hits, yet different frequencies). Assuming the Bi ion was allowed enough time to fully associate with the peptide, the method of delivery (BiBr_3_ vs gastrodenol) should not affect the outcome in theory. While gastrodenol is fully water-soluble, BiBr_3_ is not and must be added from a DMSO stock. These differences between the Bi sources may impact the screening outcomes. However, the NaAsO_2_ replicates provide additional insight, suggesting that even with the same water-soluble metalloid form, NaAsO_2_ replicates only share an R^2^ of 0.54, while the Bi conditions have an R^2^ of 0.52. Viewed in this light, it would suggest that the effects of the Bi sources are negligible, as the degree of correlation is indistinguishable from true replicates (NaAsO_2_). This highlights the importance of including a control in high-throughput screening experiments, especially if aiming to compare between conditions, as the system’s inherent variability may cause similar conditions to appear different.

### Case 5: Machine learning guided iterative design of NucB

In addition to mRNA and phage display, directed evolution experiments can also be tracked by deep sequencing over several generations. Thomas et. al., recently generated an error prone library of nuclease B (NucB) that was screened and sorted for activity (round 1) using microfluidics to construct an initial pool of functional sequences (Thomas *et al*., 2025). Subsequent rounds of iterative machine-learning based evolution were then performed to increase activity. This was achieved by training a model on the measurements of the previous round for 3 more rounds (rounds 2-4), each of which were sorted by fluorescence and sequenced (>1M reads). Here, to compare data across different rounds, we have analyzed the population at ∼90% percentile activity cutoff in each round, which is selective, but not restrictive. This study uses frequencies before and after screening to calculate enrichment (which mitigates effects of starting library bias), which we also use where appropriate.

Over the four rounds, we observe some variants exhibiting above 100-fold enrichment, while the rest are gradually less enriched over time (Fig. 6a). Despite the introduction of new variation between each round, there is still some overlap between variants of each round, with slightly higher correlation between rounds 3 and 4 compared to rounds 2 and 3 (Fig. 6b). Looking at the frequency of mutations in the selected sequences, we can see some repeated patterns between rounds that indicates the design method has already settled on many individual mutations early on (from round 2) (Fig. 6c). The mutated positions that pass sorting show a gradual shift, with more C-terminal mutations (pos 135-140) in round 2 shifting to more N-terminal mutations (pos 55-70). In terms of combined mutations in the full sequence, we can see a gradual change from a more clustered subset to a broader spread across sequence space (Fig. 6d). The spread in diversity by position (Fig. 6c) and sequence (Fig. 6d) are both consistent with the spread of designed mutations presented in the original study (Thomas *et al*., 2025). The decrease in enrichment in the rest of sequence space also highlights hotspots in sequence space that the design/selection process is gradually moving the population towards.

**Figure 6.**
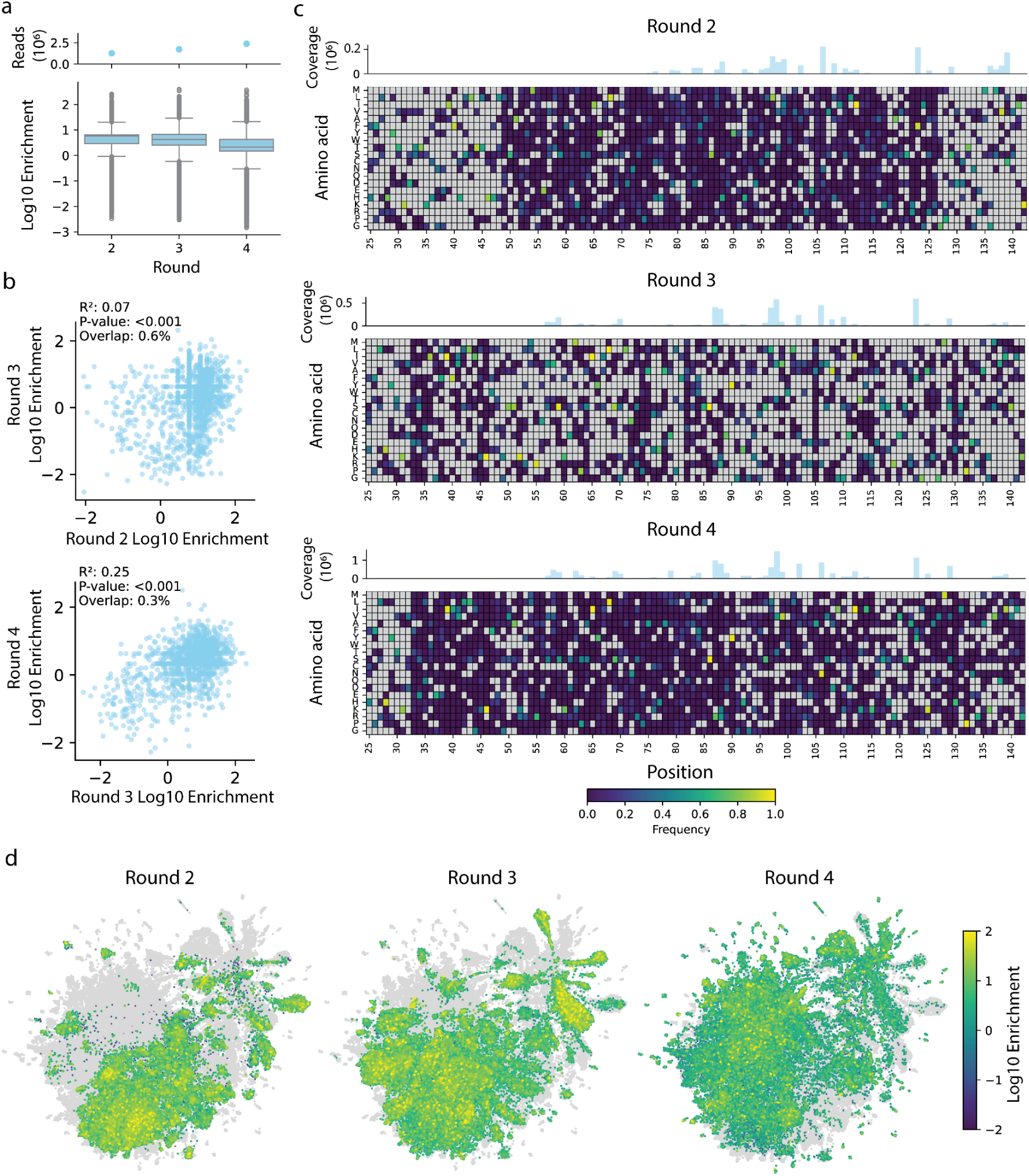
Selection in NucB iterative design libraries. (**a**) Total number of reads and sequence enrichment after each round of design and selection. (**b**) Correlation of sequence frequencies between consecutive rounds of selection. The R2 and P-value are shown for a linear fit. The overlap between rounds is calculated as sequences shared/total unique. (**c**) Heatmap of positional frequencies for each mutated amino acid. Coverage of the position is shown at the top. (**d**) UMAP display of sequences over the course of selection. All encountered sequences are in grey, while sequences present in the current round are colored and have point sizes scaled by frequency.

In this study, there is a shift in library composition between rounds due to the introduction of designed mutations based on the sequences of the previous round. The analysis here reflects both the available mutational diversity and the decisions of the design model. This is an interesting phenomenon that is not commonly analyzed, and we can infer the dynamics in the round to round correlation plots (Fig. 6b). The small overlap of just 0.3-0.6% of sequences between consecutive rounds emphasizes the change in library composition between rounds. From round 2 to 3, the vast majority of shared sequences have enrichment > 0 in round 2 (x-axis), and it is from this enriched pool that further enrichment and depletion occur in round 3 (y-axis). From round 3 to 4, the selection seems to stabilize as variants produced in round 3 show less depletion going into round 4 and the density of most shared variants are above enrichment of 0 for both rounds. Hence, despite the added mutational diversity between rounds, the methods used for other datasets are still helpful in dissecting the selection dynamics.

### Case 6: Directed evolution of an intracellular nanobody

Finally, we investigated an *in vivo* selection-evolution experiment by Cole et. al., who have pioneered a platform for directed evolution in mammalian cells using virus-like vesicles to propagate the target protein(Cole *et al*., 2025). One of the target proteins was the p53-interacting nanobody Nb139, which was evolved to select for enhanced intracellular function as a biosensor. Up to 34 rounds of growth and selection were performed, with mutations introduced in each round by the alphavirus error-prone RNA-dependent RNA polymerases. Samples were taken roughly every 5 rounds and analyzed for deep sequencing to identify mutations in a single mutation variant call format. The median read depth by position was in the order of ∼100k (Fig. 7a).

**Figure 7.**
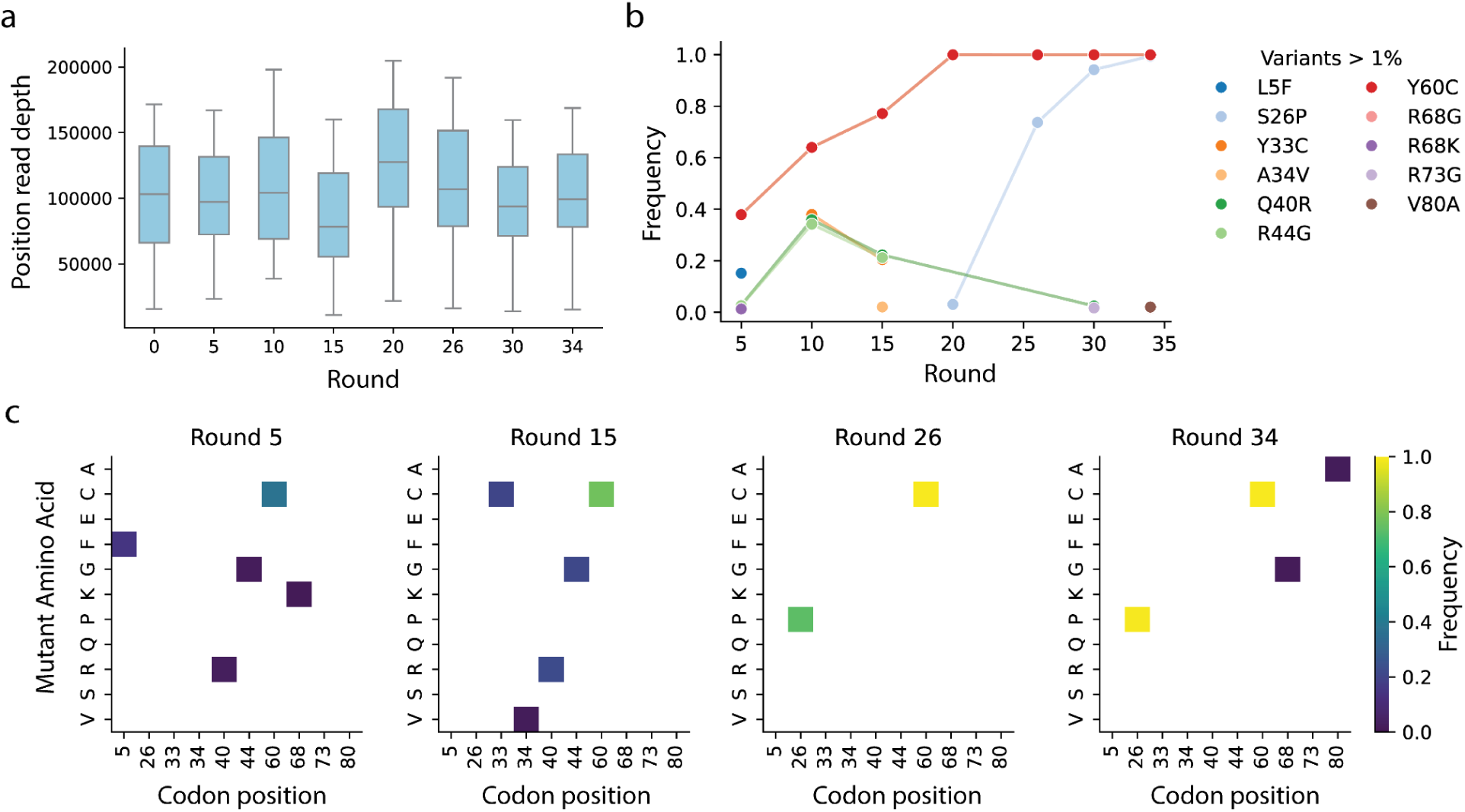
Selection outcomes in nb139 directed evolution. (**a**) Distribution of sequencing read depth by DNA position after each round of selection. (**b**) Frequencies of mutations >1% frequency over the course of selection. (**c**) Heatmap of positional frequencies for each mutated amino acid >1% frequency.

Compared to other experiments, there were far fewer mutations detected in each round, which is likely a consequence of lower starting diversity (in this experiment the evolution began from a single wild-type sequence, rather than a pre-mutated pool) and potentially higher selection pressure (selection is survival based, rather than binding/fluorescence based). The original study defined 0.3% frequency as the detection limit for mutations; here, we use a more conservative 1.0% to focus on the major mutations (Fig. 7b, c). As observed in the original study, mutations Y60C and S26P eventually dominate the population, and both mutations enhance biosensor sensitivity. Interestingly, a triple mutant of A34V/Q40R/R44G also became prominent at round 10 (∼40% frequency) but was overtaken by Y60C in later rounds. This suggests other mutational solutions to improve the desired function exist, but may not be explored due to low mutation rates, or would have been out competed due to high selection pressures. This is further underscored by the fact that S26P mutation had a greater effect than Y60C in just the wild-type background, but arose much later in the experiment due to the stochasticity of mutation accumulation (Cole *et al*., 2025).

There are signs that the sequence space covered by this directed evolution experiment is much more constrained than other case studies using large, pre-mutated libraries, and modifications to the selection strategy could have potentially broadened the solution space. The bottleneck appears to be the rate at which the error-prone RNA polymerase introduces mutations between rounds, combined with a high selection pressure that quickly fixes variants (Fig. 7b). Starting with a diverse library of the WT and a lower selection pressure (if it can be controlled) in early rounds would allow more sequence space to be explored (Yang *et al*., 2019; Erdoğan *et al*., 2023). An increase in selection pressure at later rounds can then be applied to the higher diversity pool of selection candidates to finalize the selection. Such considerations can of course be extended to systems outside of this specific study, where the starting diversity is much lower than a synthetic or naturally sampled library.

## Discussion

In trying to build a global understanding of multi-round selection experiments, we constructed a toolset for the analysis of highly diverse, large-scale deep sequencing data. The methods are broadly applicable to data from different systems and experimental designs and can help to understand selection dynamics. When applied to studies with differing experimental conditions, the toolset helps to identify common trends and differences to each system. For example, successful selection is usually accompanied by shifting population frequencies. The round-to-round correlation reveals the noise in the system and if the top hits are being consistently selected for in subsequent rounds. The amino acid preference and biophysical properties can highlight the enriched sequence patterns and provide an intuitive check for the strength of selection. The sequence space visualization provides an intuitive overview of the shifting library, highlighting changes and hotspots throughout selection. While we mainly utilize the toolset to perform post-hoc analyses, we believe it can be helpful for preliminary analysis of ongoing selection experiments. Coupled with the case studies here, the toolset can help gauge the effectiveness of selection, the degree of diversity and the resulting protein or peptide patterns from round to round. The comparison of different datasets has also led to some considerations for successful selection experiments.

### Diversity and coverage in library design

Screening large synthetic or natural libraries can circumvent gaps in engineering knowledge by drawing from a wide pool of molecular solutions. Success depends on sufficient representation of sequence diversity, as even in libraries ranging from 10^6^-10^12^ in size the number of hits can still be rare. One challenge is the coverage of the library used in the experiment, particularly for synthetic randomized libraries where the possible variation exceeds the screening capacity. Even with mRNA displays, screening ∼10^12^ variants would mean a 15-mer library still receives sparse coverage (∼10^-8^, or 1 in 100 million coverage); Repeated sampling would explore completely different sequences. While such experiments do yield hits, the results are likely subject to chance and may miss more optimal sequences (Sieber *et al*., 2015). Unfortunately, the current boundaries on library size remain due to technical limitations, and so an alternate strategy may be designing more focused libraries. While random libraries have the power to explore sequences unseen in nature, leveraging naturally existing functions (Dietrich *et al*., 2022; Cole *et al*., 2025), or semi-rational designs based on previous screens (Yang *et al*., 2019; Dobbelaere *et al*., 2021; Thomas *et al*., 2025) can drastically reduce the search space. Directed evolution and nanobody/antibody panning leverage natural selection for pre-existing binding/enzymatic activity before further optimizing for a target property. Meanwhile, iterative, semi-rational designed libraries (NucB) can traverse the sequence space step-wise without exceeding the screening capacity. However, starting from a few or just a single sequence (e.g., the WT) could lead to lower accumulation of diversity that may miss possible solutions. In such cases, starting with a larger sequence pool, increasing the mutation rate, or reducing the selection pressure to allow some genetic drift may help explore a broader range of possible solutions (Yang *et al*., 2019; Erdoğan *et al*., 2023).

### Effects of selection systems

The accessibility of hits is also dependent on the protein system of choice and method of selection. A library of variants, especially a fully randomized library, has the potential to contain any number of unknown functions and could end up enriching for unintended products if not guided towards the correct target. Ideally, the selection system would be able to directly select for the property of interest (e.g., activity, inhibition) (Nikoomanzar *et al*., 2019; Starr *et al*., 2020; Chen *et al*., 2022), but in most cases a proxy property (e.g., binding instead of inhibition) has to be used instead due to technical limitations. For binding experiments, one should ensure that the target is in the intended form; in the case of M^pro^, the presence of excess monomers may have diluted targets for the functional dimer (Johansen-Leete *et al*., 2022). If the target protein is large, focusing on the smallest unit with the intended target region may help reduce unwanted binding surfaces. Furthermore, the selection strength needs to be adequate to match library diversity, allowing rare functional variants to better compete with the high background diversity and reducing the time to enrich hits. Strong selection is particularly important in early rounds, where low initial frequencies for potential hits make them vulnerable to loss by genetic drift (Otto and Whitlock, 2013; Martínez and Lang, 2023). Ideally, the system allows for direct control of selection pressures, such as drug concentration for cell-survival screens (Chen *et al*., 2022; Erdoğan *et al*., 2023), or signal gated droplet/cell sorting (Klesmith *et al*., 2017; Starr *et al*., 2020; Thomas *et al*., 2025). The addition of an existing competitor may help increase selection for stronger binders (e.g., streptavidin in the bismuth peptides experiment) (Ullrich *et al*., 2024), or more rigorous negative selection against non-binders could be employed (reducing bead binders in the spike and M^pro^ peptide experiments) (Thijssen *et al*., 2023). Although often less malleable than the library design, the selection system still warrants scrutiny when troubleshooting and optimizing selection experiments.

### Potential uses of large-scale selection datasets in machine learning

High-throughput sequencing on high diversity libraries has great value as knowledge bases for machine learning based design of proteins and peptides (Dobbelaere *et al*., 2021; Kouba *et al*., 2023). Generally, machine learning of proteins trains models to build (often complex) correlations of patterns in sequence or structure to a set of phenotypes (activity, affinity, etc.), and using the learned correlations to further predict function of unknown sequences or generate novel functional sequences. The quality of the measured sequence data and the phenotypes greatly affects the outcome of machine learning, and noise or lack of coverage would hamper performance (Kouba *et al*., 2023). The sequencing data should be of sufficient quality and depth to fully represent the diversity and functionality of the library, so that the correct patterns are represented during training. Furthermore, the coverage of the library should be dense enough to provide adequate numbers of examples for each group of functional sequences; if the functional sequence space is only sparsely sampled, there would be little chance to establish a learnable pattern (Dobbelaere *et al*., 2021). However, screening data metrics (frequency, enrichment) can be noisy in practice, and may not reflect their performance in low-throughput biochemical assays (Johansen-Leete *et al*., 2022). In the case of noise, negative controls and characterization of a subset of hits should be used to establish baselines that distinguish function from noise. High throughput screening has the potential to generate large, labelled datasets that can be harnessed to enhance machine learning based protein/peptide design (Yang *et al*., 2019; Minot and Reddy, 2024; Thomas *et al*., 2025), and this potential should be leveraged by carefully exploring and cataloging the functional sequence space.

## Methods

### Datasets

Data for the SARS-CoV-2 main protease and *Plasmodium falciparum* Pfs230 selection were obtained as raw fastQ files from the corresponding authors. The SARS-CoV-2 spike dataset (also fastQ format) can be publicly accessed (https://doi.org/10.34894/WJRHLK). For raw fastQ files, the coding portion was determined, and filtered to be above 20 Phred score for all positions before converting from DNA to amino acid sequence. For the two SARS-CoV-2 mRNA displays, the fixed starting initiator (D- or L-Tyr) and linkers were trimmed from the sequence before analysis, leaving just the randomized region. The nanobody sequences from phage display were analyzed using IgBlast with a custom alpaca database built from the IMGT alpaca IGHV references to identify the CDR3 region (Manso *et al*., 2022). Data for the bismuth bicyclic peptides and iterative design of NucB were available as precompiled counts in the supplementary data and were used without additional modification. Data for the directed evolution of nb139 was obtained in variant call format (vcf) from the author’s provided Gene Expression Omnibus entry (GSE250502).

### General analysis methods

We demonstrate the use of a series of analyses to examine various aspects of the selection process. We note that the methods are not a set of strict protocols, but rather a series of concepts to be adapted to the dataset as applicable. The application of methods on each dataset is detailed in the accompanying coding example (https://github.com/johnchen93/SelectionDataAnalysis). The general method is broadly described below.

To gain an overview of sampled diversity and frequencies. We examine the total sequencing read count per sample. This can inform the user if the coverage is sufficient to analyze certain sequence frequencies. For example, a frequency of 0.1% would be reliable with 10^6^ total reads (100 variant reads), but is likely noise with 10^3^ total reads (1 variant read only). In the case of variant call format (VCF) we used the per DNA position read depth.

Shifts in population diversity are plotted as distributions across rounds, to assess shifts across selection. When selection acts to enrich certain subpopulations, we expect either an increase in the upper frequencies and/or a drop in lower frequencies as a result of separation between those selected for and against the desired function. When there is sufficient non-selected data (such as round 0 in the NucB dataset), enrichment ( selected frequency / non-selected frequency) can be analyzed in a similar fashion.

To examine consistency between successive rounds, we use a scatterplot of frequencies. Higher correlation indicates that successive rounds share similar patterns of high vs low frequency sequences. As the population changes over the course of selection, we do not expect perfect correlation in most cases. The plots themselves can be a quick visual check to see that top hits in one round are consistently carried to the next (linear or monotonic). As the libraries typically carry a large diversity of non-hits, the dispersion at lower frequencies is expected, and the spread would correlate with the degree of noise in the experimental system.

For viewing sequence patterns, we apply both a heatmap for single amino acid frequencies, as well as a biophysical line plot to examine patterns across whole sequences. The heatmaps are plotted from the position weight matrices of the amino acids (frequency of each position and amino acid, disregarding linkage to other positions) of aligned sequences (in practice, no alignments are necessary except for variable length regions such as the CDR3 of antibody/nanobodies). The biophysical line plots depict the top 100 sequences in terms of frequency, with each position represented by the hydrophobicity (Kyte and Doolittle)(Kyte and Doolittle, 1982) or the accessible surface area (absolute ASA calculated by PyMol, normalized to the Miller 1987 reference set)(Miller *et al*., 1987) of the amino acid. Lines connect entire sequences, and overlapping regions highlight shared patterns. The average and envelope (standard deviation) are also plotted to gauge the mean and spread in positional patterns. The heatmaps are an intuitive view of the abundance of amino acids (or if a position is varied at all), while the biophysical line plots can help reveal patterns across several positions.

To obtain a comprehensive overview of sequence diversity over the course of selection, we use the pLM Evolutionary Scale Modelling (ESM, version 1b) (Rives *et al*., 2021) (https://github.com/facebookresearch/esm), to embed protein sequences into mean-pooled numerical vectors. The vectors (1280 features in length) are reduced to 30 features by PCA, before applying dimensionality reduction via uniform manifold approximation and projection (UMAP)(McInnes *et al*., 2020) to 2 dimensions. The sequences are then plotted as scatter plots by round, with color indicating the frequency (or enrichment). Absolute distances in the UMAP plot have no meaning, but sequences that are further away indicate they are different, while those closer to each other are more similar. The UMAP is useful for a complete and intuitive overview of the shifting of sequence populations across rounds, as well as showing any shared hot spots.

## Acknowledgements

This work was funded by the Australian Research Council Center of Excellence in Synthetic Biology (grant CE200100029) and the Australian Research Council Center for Innovation in Peptide and Protein Science (grant CE2001 00012). W-H.T. is supported by National Health and Medical Research Council of Australia (NHMRC) (grants GNT2016908 and APP2001385). C.N. is a recipient of the Australian Research Council Future Fellowship (FT220100010). J.Z.C. is a Human Frontier Science Program postdoctoral fellow (LT0048/2023-L).

## Supplementary Data

**Supplementary figure 1.**
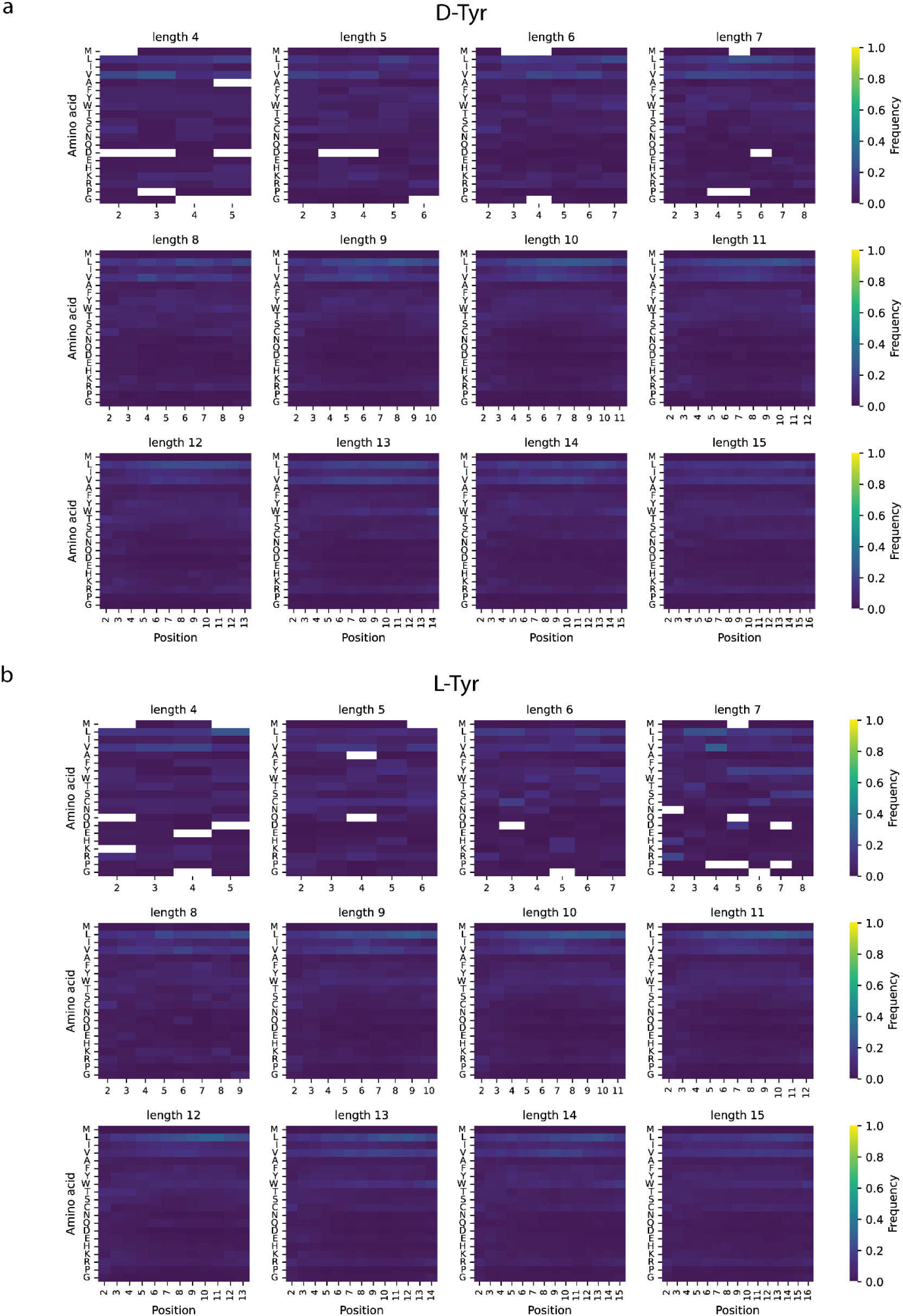
Per position amino acid frequency of MPro peptide screen at round 9 for all lengths. (a) Heatmaps for D-Tyr library. (b) Heatmaps for L-Tyr library.

